# Unexpected finding of *Fusobacterium varium* abundance in cattle rumen: implications for liver abscess interventions

**DOI:** 10.1101/2022.12.05.519234

**Authors:** Cory Schwarz, Jacques Mathieu, Jenny Laverde Gomez, Megan R. Miller, Marina Tikhonova, T.G. Nagaraja, Pedro J.J. Alvarez

**Author notes:** Address correspondence to Jacques Mathieu.

## Abstract

*Fusobacterium varium* has been generally overlooked in cattle rumen microbiome studies relative to the presumably more abundant, liver abscess-causing *Fusobacterium necrophorum*. Here, we revisit that conventional wisdom and report greater relative abundance of *F. varium* than *F. necrophorum* in both raw rumen samples and in lactate-supplemented enrichments tailored for *F. necrophorum* growth, despite its consistent inadvertence in past ruminal surveys and putative inability to metabolize lactate. Our observation that *F. varium* grows under restrictive conditions used to enumerate *F. necrophorum* suggests that previous estimations were inaccurate and *F. varium* is an underestimated player within the ruminal community. Exposure to tylosin, the current gold standard among prophylactic liver abscess prevention strategies in cattle, consistently reduced growth of all *F. necrophorum* strains screened by greater than 67% relative to unexposed controls. In contrast, *F. varium* strains were completely or highly resistant (0 - 11% reduction in maximum yield). Monensin, an ionophore fed to cattle to improve feed efficiency also had stronger inhibitory activity against *F. necrophorum* than against *F. varium*. Finally, preliminary genomic analysis of two *F. varium* bovine isolates revealed the presence of virulence genes related to those of pathogenic *F. varium* human isolates associated with active invasion of mammalian cells.

**Importance:** Judicious antibiotic use is essential to mitigate the spread of antimicrobial resistance. Dogmatic prophylactic use of in-feed tylosin to control cattle liver abscesses hinges on the assumption that *F. necrophorum* in the rumen is the main etiologic agent. However, our unexpected finding of abundance of *F. varium* in the rumen and its resistance to antibiotics, in hand with the potential pathogenicity of this species, calls for increased attention to *F. varium*. Further investigation into *F. varium* is necessary to better understand bovine liver abscess development and devise higher-precision alternatives to antibiotic treatment.

## INTRODUCTION

*Fusobacterium* is a genus of gram-negative non-spore-forming nonmotile anaerobic bacteria commonly found in human and animal gastrointestinal tracts. Several species are known to be pathogenic, including *F. nucleatum* and *F. necrophorum*, which have been extensively researched due to their association with a wide range of infections and diseases [1-4]. Within the feedlot beef cattle industry, *F. necrophorum* is considered the primary etiological agent of liver abscess development [5, 6], which cause significant financial impact due to liver condemnation, decreased animal productivity and carcass value [7]. Currently, antibiotic feed additives that inhibit the growth of *F. necrophorum*, such as tylosin and virginiamycin, are the most common method for preventing liver abscesses (LA) [8]. However, there is substantial public and regulatory pressure to reduce overall antibiotic use to curtail proliferation of antibiotic resistant bacteria. As such, other methods are being explored to prevent LA formation in cattle, including vaccination [9], dietary supplementation with phytogenic additives (e.g., essential oils) [10, 11] or *Saccharomyces cerevisiae* fermentation products [12]. However, such approaches have only proven effective *in vitro* or under strictly controlled conditions and have not been widely adopted within the cattle industry [13, 14].

As an alternative to conventional antibiotics for liver abscess prevention there is growing interest in more selective microbial control technologies that can facilitate precise reductions in the concentrations of target pathogens (e.g., *F. necrophorum*) and minimize potential unintended impacts. In this regard, there has been increasing academic and industrial interest in the use of bacteriophages (phages) as animal feed additives for biocontrol and microbiome editing applications [15, 16]. In efforts to develop phage cocktails for several bovine ruminal species, including *F. necrophorum*, we surprisingly found that a widely overlooked species of *Fusobacterium* – *Fusobacterium varium –* was predominant in bovine rumen samples obtained from different animals and disparate geographical locations. The only prior mention of *F. varium* in cattle rumen was over 40 years ago in a rarely cited study [17] conducted in Japan that described it as being the most commonly isolated *Fusobacterium* species from ruminal fluid (at more than 10^5^ cfu/mL), and there is little further mention of this species within the field of ruminal microbiology. The lack of information and characterization of ruminal *F. varium* led us to investigate the identity and distribution of *Fusobacterium* species in the bovine rumen.

This paper presents converging lines of evidence from targeted metagenomic and culture-dependent investigations that *F. varium* is both more prevalent and abundant than *F. necrophorum* in cattle rumen, contrary to the current dogma. This finding is important because previous studies that relied on most probable number (MPN) assays [18-20] may have erroneously included *F. varium* when determining *F. necrophorum* ruminal concentrations, subsequently impacting those findings, including treatment efficacy. Moreover, the vast majority of targeted metagenomic studies of the bovine rumen (i.e., 16S rRNA analyses) have been conducted using methods that provide only genus-level taxonomic resolution. As such, it is currently unclear whether *F. varium*, an emerging human pathogen [21], plays a role in bovine LA etiology or other diseases. Thus, a better understanding of the distribution and role of *F. varium* in the bovine rumen is necessary for the development of selective biocontrol technologies for LA prevention.

## MATERIALS AND METHODS

### Rumen fluid sampling and storage

Rumen fluid samples were obtained from several different locations within the United States over a three year period. In February 2018, fifty samples were taken immediately post-slaughter from a USDA-monitored cattle processing facility in Amarillo, Texas. The samples collected included cattle with apparently healthy livers or abscessed livers and were taken directly from harvested rumen by filling 50 mL centrifuge tubes (no headspace) and immediately placing on ice. Oxyrase (10% v/v) was added to 10 tubes to assess any potential benefit for maintaining anaerobe viability during sampling and transport. The samples originated from five different feedlots within Texas, with ten different cattle sampled from each site. All samples were shipped overnight on ice, processed within 24 h, diluted 1:1 with a 20% DMSO solution, and stored at -80°C until further use. In 2020 and 2021, another 100 rumen fluid samples were provided by Kansas State University (Prof. T.G. Nagaraja), with 10 of these sample taken from cannulated cattle housed at their veterinary facility and 90 obtained from three separate feedlots as part of a liver abscess study. Approximately half of these samples were from animals with liver abscesses. These samples were shipped overnight on ice, but contained some headspace in most cases. In addition, 15 rumen fluid samples were obtained from the USDA Livestock Issue Research Unit (Lubbock, TX) in 2021, with 7 from cattle that did not receive tylosin in their rations. Besides these cattle, the diet and antibiotic status of the cattle were not known. The samples from KSU and the USDA were cryopreserved in the same manner as those from the processing facility. All samples used in this study originated from feedlot cattle within Texas, Nebraska, Kansas, or Missouri.

### Bacterial strains and culture conditions

*Fusobacterium varium* strain designation NCTC 10560 (ATCC 8501™), and *Fusobacterium necrophorum* subspecies *necrophorum* strain VPI 2891 (ATCC 25286™) were obtained from the American Type Culture Collection (ATCC, Manassas, Virginia, United States). Previously isolated *F. necrophorum* strains were kindly provided by Prof. T.G. Nagaraja (KSU), including 8 from subspecies *necrophorum* and 5 from subspecies *funduliforme*. All *Fusobacterium* strains were grown under anaerobic conditions in Hungate tubes (Chemglass Life Sciences, Cineland, NJ, United States) in Brain Heart Infusion (BHI) broth (Teknova, Hollister, CA, United States) supplemented with 1 mg/L of resazurin and 5 g/L of both peptone and yeast extract, and incubated at 37°C. Additional broth media used in these experiments included anaerobic peptone yeast (PY) broth (Anaerobe Systems); Medium II, a semi-defined medium [22]; and a modified lactate medium (MLM) [20].

### Isolation of *Fusobacterium* species on selective medium by direct plating

Ten microliters of ruminal fluid or ruminal contents suspended in PBS were directly streaked onto selective agar plates [23] and incubated anaerobically at 37°C up to 4 days. The selective medium included the antibiotics josamycin, vancomycin, and norfloxacin (at 3, 4 and 1 μg/ml, respectively; JVN) as selective agents, plus 5% defibrinated horse blood in fastidious anaerobe agar (FAA) (Neogen, Lansing, MI, United States). When growth was visible on the plates, individual colonies were picked, streaked, and used as template for colony polymerase chain reaction (cPCR) to identify the organism by amplification of the 16S rRNA subunit using the universal 27F (5’-AGAGTTTGATCCTGGCTCAG-3’) and 1492R 5’-GGTTACCTTGTTACGACTT-3’) primer set [24]. Following visualization, bands of the appropriate size (∼1,400 bp) were cut, gel extracted (E.Z.N.A.® Gel Extraction Kit, Omega Biotek, Norcross, GA, United States) and sent to an external lab for Sanger sequencing (Genewiz from Azenta Life Science, South Plainfield, NJ, United States).

### Selective enrichment of ruminal content samples

To increase the abundance of *Fusobacterium* species for isolation, anaerobic culture medium PY was amended with 50 mM lactate with or without JVN antibiotics (added after autoclaving). Briefly, 1 g of rumen digesta was suspended in 10 mL of PY medium and 100 µl aliquots were added by syringe to pre-sterilized Hungate tubes containing anaerobic PY medium amended with JVN and lactate. These tubes were incubated overnight at 37 ° C. Subsequently enrichments were plated on FAA-JVN medium for isolation of *Fusobacterium* species or used for 16S rRNA amplicon sequencing and bacterial community analysis.

### Genomic DNA extraction and quantification

Genomic DNA of enrichment communities for use in PCR was extracted using the Qiagen DNA Stool Kit, using cell pellets of the enrichment instead of stool sample, but otherwise according to the manufacturer’s instructions. Samples were further purified using a Zymo Clean & Concentrate Kit prior to PCR. Genomic DNA of bacterial isolates for genome sequencing was extracted via a slight modification of a published method [25] to optimize for long DNA fragments. Briefly, 2 mL of logarithmic growth phase culture was pelleted by centrifugation and resuspended by vortexing in B1 buffer (Qiagen, Hilden Germany: 50 mM Tris • Cl pH 8.0; 50 mM EDTA pH 8.0; 0.5% Tween 20; 0.5% Triton-X100) amended with 2 µL of RNaseA solution (100 mg/mL) and 100 µL of lysozyme solution (100mg / mL). After an hour of incubation at 37°C, 45 uL of Qiagen proteinase K stock solution was added and incubated at 50° C for 1 hour. Then, a 1:1 volume of biotechnology-grade phenol/chloroform/isoamyl alcohol was added and agitated by hand. Phase separation was achieved by centrifugation at 3,000 x g and the upper aqueous layer was transferred by pipetting to a new sterile microcentrifuge tube. This step was repeated a total of three times, followed by a final cleaning step using a 1:1 volume of chloroform/isoamyl alcohol 24:1 to the aqueous sample, followed by centrifugation and aspiration and retention of the upper aqueous layer. This was followed by ethanol precipitation [26] and resuspension of the DNA in 100 uL of molecular grade water. Quantification of concentration and quality of DNA extractions were performed using a NanoDrop DN-1000 Spectrophotometer (Thermo Fisher Scientific, Waltham, Massachusetts, United States).

### Genome sequencing and assembly

Libraries for genome sequencing were prepared using high molecular weight DNA prepared by phenol-chloroform extraction (detailed above) using a Ligation Sequencing Kit (SQK-LSK009, Oxford Nanopore Technologies) and sequenced on a MinION sequencer (Oxford Nanopore Technologies, Oxford, United Kingdom) using a Flongle adapter and flow cell (R.9.4 chemistry) according to the manufacturer’s instructions. MinKNOW software was used to basecall and demultiplex reads. Reads were then trimmed using porechop and those less than 1000 bp were filtered out using seqkit. Trimmed and filtered reads were then assembled using the miniasm + Racon pipeline, as previously reported [27]. Assembled contigs were then uploaded to the RAST [28-30] server for automated gene calling and functional annotation.

### 16S rRNA amplicon sequencing and community analysis of raw ruminal content samples and enrichments

16S rRNA subunit amplification was performed using 1 µL of diluted (10%) rumen fluid or enrichment culture, or gDNA extracted as described above, directly as template for PCR using KAPA HiFi Hotstart ReadyMix (Roche, Basel, Switzerland). Sequencing libraries were prepared using the Oxford Nanopore 16S Barcoding Kit (SQK-RAB204) according to the manufacturer’s instructions. Libraries were sequenced using an Oxford Nanopore MinION sequencer with a Flongle adapter and flow cell, and reads were basecalled and demultiplexed using MinKNOW. Processed reads were then classified using either blastn against a local copy of the NCBI 16S rRNA database, or Emu using the provided database [31].

### *Fusobacterium* genus-specific primer set amplification and sequencing

A region of the 16S rRNA subunit was amplified using previously reported *Fusobacterium*-specific primers FUSO1 (5’-GAGAGAGCTTTGCGTCC-3’) and FUSO2 (5’-GGGCGCTGAGGTTCGAC-3’) [32]. An annealing temperature of 60°C was used, producing amplicons of approximately 610 bp. These amplicons were then sequenced using the Oxford Nanopore Ligation Sequencing Kit according to the manufacturer’s instructions and analyzed in the same manner as the 16S rRNA workflow described above.

### Antibiotic sensitivity assays

Mid-log bacterial cultures were adjusted to an OD_600_ =0.1 using fresh medium and added to a 96-well plate via multichannel pipette. Antibiotics were added to medium supplemented with 0.1% freshly prepared cysteine-HCl [33] and 5% Oxyrase, plates were then sealed using qPCR grade plate films to prevent gas exchange, and incubated at 37° C. All growth curve data were collected in triplicate in at least two separate experiments on a Tecan Infinite M Plex (Tecan Group Ltd., Männedorf, Switzerland) at a minimum one hour sampling interval for at least 15 hours. Due to the different growth rates between organisms and short stationary phases, comparisons were performed using each strain’s growth maxima, identified by highest respective OD_600_. The representative concentrations selected for the tylosin tartrate (45 mg/L) and monensin sodium sulfate (3.5 mg/L) were determined from the literature [34-36]; briefly, concentrations were selected based on the recommended amounts of each antibiotic fed per cattle head per day, assuming a ruminal compartment volume of 100 L and twice daily feedings. A minimum of three biological and technical replicates were used for each study (e.g., at least three reactions of three different bacterial strains, or nine in total).

### Phage isolation and host range testing

Bacteriophage pools were prepared from lactate-amended enrichment cultures inoculated with ruminal fluid. Briefly, enrichment cultures were harvested after overnight incubation and centrifuged in 15 mL tubes at 10,000 RCF for several minutes to remove cell debris. The supernatant was then aspirated and filtered into sterile 15 mL centrifuge tubes using a 0.22 μm PVDF syringe filter. Polyethylene glycol 8000 (PEG 8000) and NaCl were then added to a final concentration of 10% (w/v) and 1 M, respectively, and the entire solution incubated at 4°C overnight to permit precipitation. Tubes were then centrifuged for 30 mins at 10,000 RCF and 4°C. The supernatant was then removed and the pellets resuspended in a volume of SM buffer approximately 10-fold less than the original culture. Double layer overlays containing different *Fusobacterium* species and strains were used to assess phage host ranges by spot assay. Briefly, 4.25 mL BHI top agar (0.4% agarose, 5% glycerol, 1 mM CaCl_2_, and 1 mM MgCl_2_) was melted by microwaving and equilibrated to 50°C in a heat block. The volume was then brought to 5 mL by addition of 250 μL of log-phase cultures, 5% lysed horse blood and 5% Oxyrase (final concentration) immediately prior to pouring on BHI bottom agar (1.5% agarose, 5% glycerol, 1 mM CaCl_2_, and 1 mM MgCl_2_). Plates were allowed to solidify at room temperature under ambient conditions for 5 mins, then spotted with 5 μL of each phage pool. Up to 20 pools were spotted on each plate, and allowed to dry for approximately 5 – 10 mins prior to incubation for 3 – 4 days at 37°C under anaerobic conditions. Plates were checked daily for evidence of lysis. Where lysis was observed, 10-fold serial dilutions of the original phage pool were prepared and spotted again on the same strain to estimate phage concentration. Individual plaques were picked, resuspended in 100 μL SM buffer, and subject to two additional rounds of purification prior to amplification on the isolation host in liquid broth.

## RESULTS

### *Fusobacterium varium* is ubiquitous and a predominant member of its genus in cattle rumen samples

During attempts to isolate ruminal strains of *F. necrophorum* using both *Fusobacterium*-selective media [23] and lactate enrichment, which is commonly used to selectively grow *F. necrophorum* [6, 20, 37], we identified the majority of isolates as *F. varium* via 16S rRNA sequencing. Out of 165 ruminal fluid samples obtained, over half have been screened, with *F. varium* being found in 100%. The processing plant samples were obtained from feedlot cattle, though no information as to their specific diet, antibiotic exposure, or liver abscess status was provided. However, the ruminal fluid samples obtained from KSU and the USDA had additional metadata describing their liver abscess status and tylosin exposure. All 15 USDA samples and at least half of the 100 KSU samples were included in those screened for *Fusobacterium* and we did not observe differences in the presence of *F. varium* due to the use of tylosin or between abscessed and non-abscessed cattle. When streaking ruminal fluid directly onto *Fusobacterium*-selective medium (JVN), approximately 50% of the strains were *Fusobacterium*, with 50 - 75% of those isolates identified by 16S rRNA sequencing as *F. varium*, and the remainder generally being *F. necrophorum* and *F. gastrosuis*. When colony picking, the frequency of *F. varium* isolates decreased over time due to improvements in visually distinguishing amongst colonies. However, when plating lactate enrichments onto JVN, we often only identified *F. varium* when randomly selecting 8 – 10 colonies for cPCR. Additionally, we observed large numbers of *F. varium*-specific phages from enrichment cultures containing lactate. We isolated and purified these phages prior to assessing their host ranges on a library of different *Fusobacterium* species. The *F. varium* phages isolated to date have proven to be species-specific, and incapable of infecting any *F. necrophorum* strains. Notably, isolating *F. necrophorum* phages from cattle ruminal fluid requires much more effort and the phages to date have all been temperate, while the *F. varium* phages appear to be both lytic and temperate, as well as present in all samples tested.

To determine the relative abundances of different *Fusobacterium* species in ruminal fluid and in various enrichment cultures, we utilized targeted 16S metagenomics to assess community composition and structure. To obtain species-level resolution, 16S rRNA amplicons were generated using a close to full-length, barcoded primer set (27F/1492R) and sequenced with an Oxford Nanopore MinION system. Community analysis confirmed the presence of *Fusobacterium* within unenriched, preserved bovine ruminal samples, with the genus-level characterization being consistent with previous studies [38]. Specifically, we found *Fusobacterium* to have a relative abundance between 0.001% and 0.003% in ruminal fluid. However, such low abundances (1 *Fusobacterium* read per 50,000 total reads) made it difficult to assess species distribution. Thus, amplicons were also generated from rumen samples using genus-specific primers [39] to increase sequencing depth and assess the distribution of *Fusobacterium* species within each sample. Out of 98,575 classified reads from 12 ruminal fluid samples, 98,052 were assigned to *Fusobacterium*, highlighting the specificity of this approach. Using this method, we found *F. varium* was the most abundant member of the genus, on average comprising 46% of *Fusobacterium* species and 0.00138% of the total community (Fig.1). The next most abundant species was *F. necrophorum* (31% and 0.00093%), followed by *F. gastrosuis* (19% and 0.00057%), and the remaining 5% (0.00015% of the community) were a collection of other *Fusobacterium* species (in descending order of abundance, *F. ulcerans, F. nucleatum, F. equinum, F. simiae, F. mortiferum, F. gonidiaformans, F. periodonticum, F. russi*, and *F. canefelium*).

**Figure 1.**
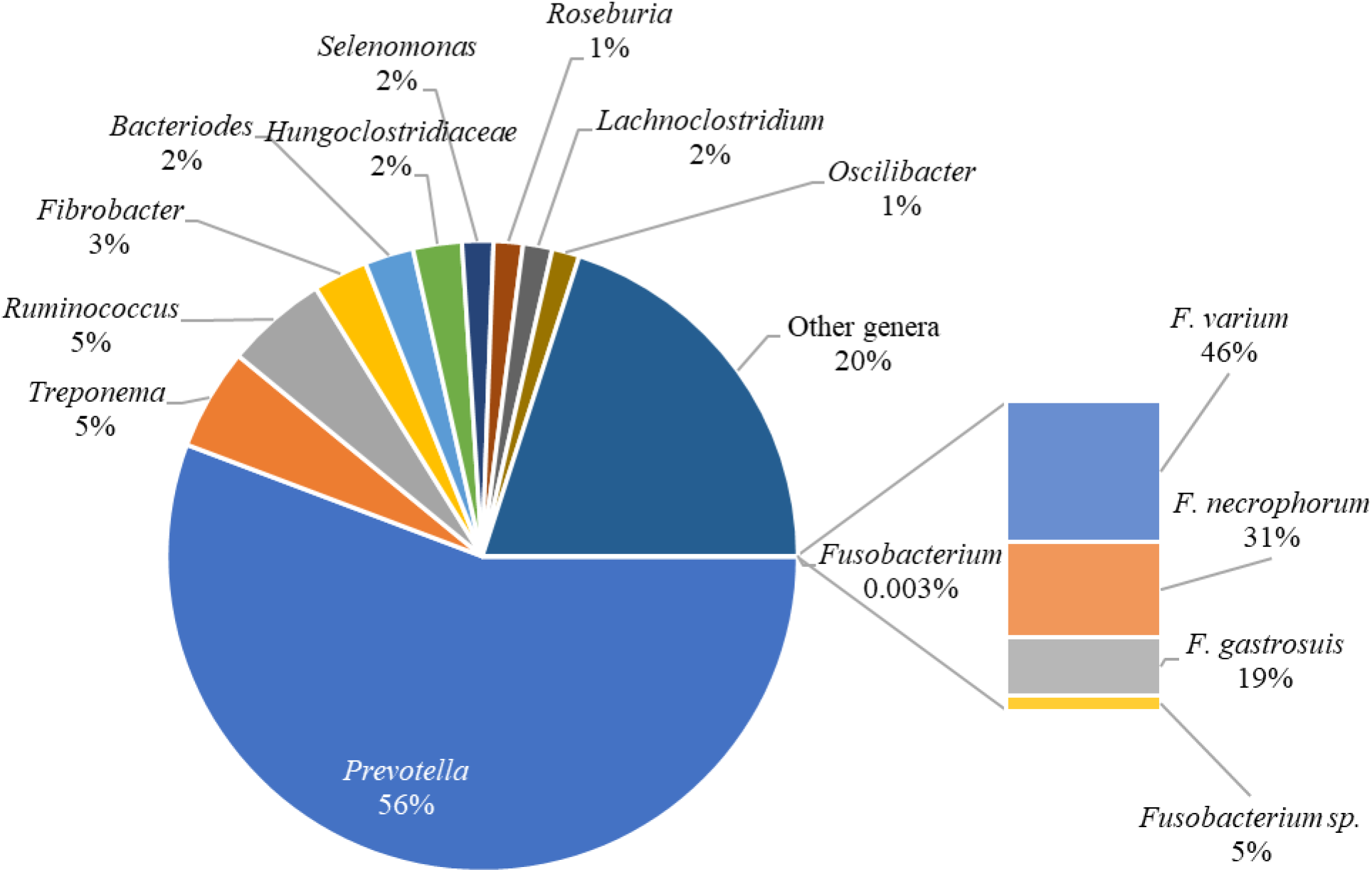
Dominance of *Fusobacterium varium* among *Fusobacterium* species in bovine rumen sample. 16S rRNA subunit sequencing and community analysis showed that the low relative abundance of *Fusobacterium* is consistent with previously published findings. However, the distribution of species within the *Fusobacterium* genus was unexpected. The abundance of each detected *Fusobacterium* species is represented by the smaller chart on the right. *Fusobacterium varium* was the most abundant species (46%) followed by *F. necrophorum* (31%) and *F. gastrosuis* (19%). Other of *Fusobacterium* species made up the remaining 5%.

### *F. varium* was unexpectedly enriched under conditions tailored to select for *F. necrophorum*

As previously mentioned, we noted an increased frequency of *F. varium* isolation after lactate enrichment. Using the same ruminal samples as above, the average relative abundance of *Fusobacterium* increased over four orders of magnitude, accounting for approximately 34% of reads (Fig. 2). The next most abundant genera were *Clostridium* (22%), *Proteus* (3%), *Paeniclostridium* (2%), *Enterococcus* (2%), *Escherichia* (1%), and *Shigella* (1%). The remaining 35% of reads belonged to genera whose individual relative abundance was lower than 1%. Within the *Fusobacterium* genus, the most prevalent species were *F. varium* (24.5% of all reads), followed by *F. ulcerans* (5.3%), *F. mortiferum* (2.6%) and other *Fusobacterium* species (in descending order of abundance, *F. nucleatum, F. gastrosuis, F. equinum, F. periodonticum, F. perfoetens, F. necrophorum, F. canifelinum*, and *F. simiae*) accounting for the final 1.1%.

**Figure 2.**
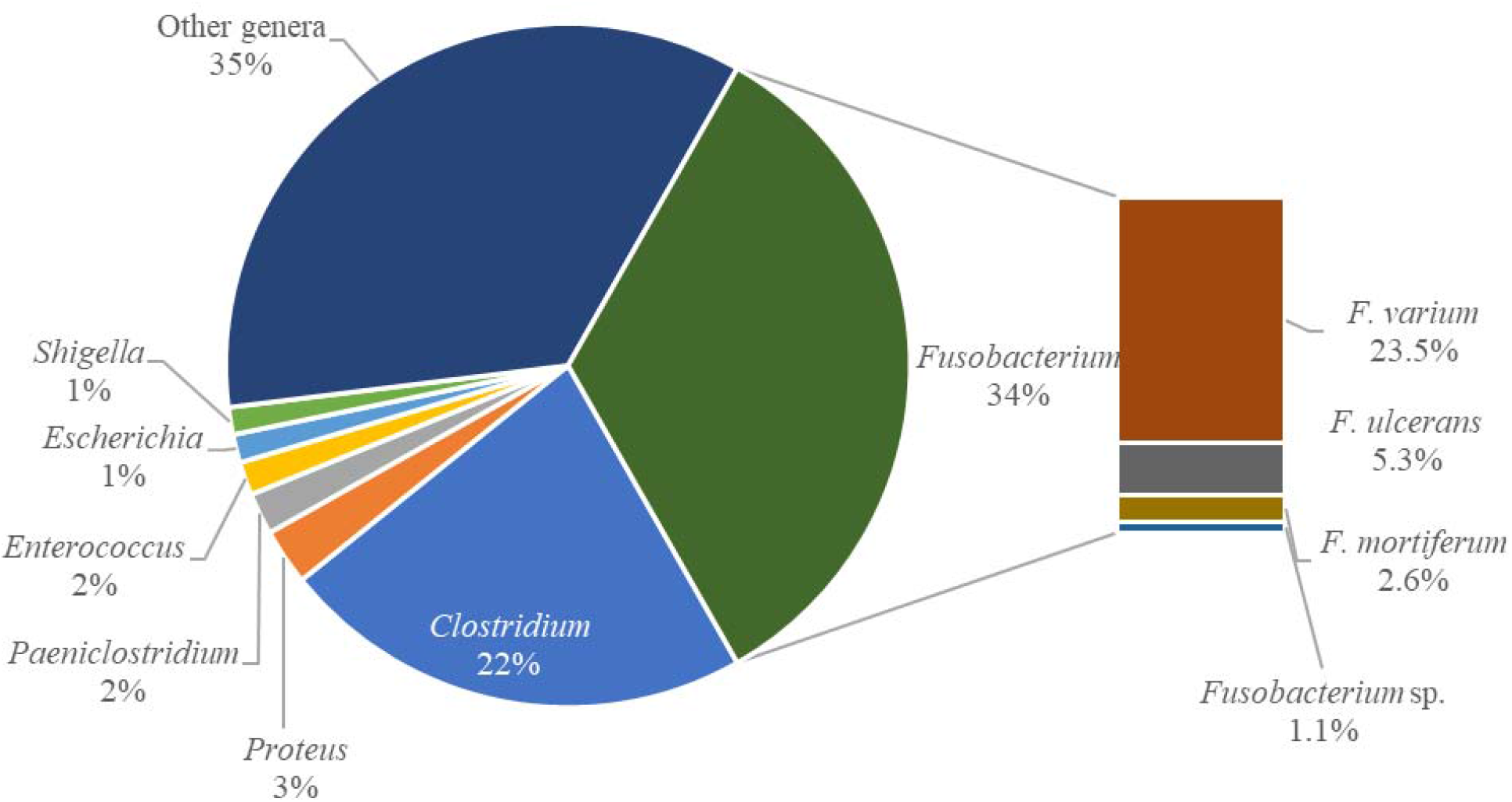
*Fusobacterium varium* was preferentially enriched under conditions designed to increase the growth of *F. necrophorum*. An enrichment of PY amended with 50 mM lactate and *Fusobacterium*-selective antibiotics (JVN) was used to increase the abundance of *F. necrophorum*. The prevalence of genera changed drastically from the ruminal sample, and though the enriched community was dominated by *Fusobacterium* (33.5% of all reads), the most prevalent was *F. varium* (24.5%).

To further understand the impact of various substrates on different *Fusobacterium* species, we prepared a duplicate series of enrichment cultures (using two different ruminal fluid samples) in anaerobic PY medium containing no amendment (controls), 50 mM lactate, 50 mM lysine, or 50 mM maltose. In addition, replicates of each were prepared with the JVN antibiotic mixture. We observed a modest enrichment of *F. varium* in controls, with an average relative abundance of 0.12%. Notably, *F. necrophorum* was not detected in either control, though each sample only averaged about 30,000 reads, so only enriched *Fusobacterium* species would be detectable under these conditions. However, the addition of lactate dramatically increased *F. varium* abundance to 71.5% while *F. necrophorum* was only slightly enriched to 0.39%. Lactate enrichments with JVN averaged 61.1% *F. varium* and 0.40% *F. necrophorum*, seemingly offering no benefit for enrichment. Interestingly, the addition of lysine appears to have negatively impacted *F. necrophorum* abundances to 0.008% and only slightly increased *F. varium* over controls to 0.32%. However, the addition of JVN increased *F. varium* abundance to 24.1% of total reads while *F. necrophorum* was 0.05%. In maltose enrichments, no *F. necrophorum* was detected while *F. varium* was 0.07% of total reads. However, JVN amendment in addition to maltose increased *F. varium* to 66.5% and *F. necrophorum* to 3.05% of total reads. Overall, these data suggest a context-dependent response to JVN antibiotics, and frequent enrichment of *F. varium* on substrates previously thought to be selective for *F. necrophorum*.

As lactic acid is believed to play a vital role in liver abscess etiology, we compared the microbial communities of several ruminal fluid samples inoculated into PY medium with and without 50 mM lactate. Six samples were selected from which we had obtained and partially characterized *F. varium* isolates. Interestingly, we observed differences in *Fusobacterium* composition and growth responses between samples. In one sample (KL10), *F. varium* was the most abundant species even without the addition of lactate, comprising over 30% of total reads, but was not further enriched by lactate. In other samples (1703, USDA-12), *F. varium* was only minimally enriched by lactate. We also noted a steady decrease in the magnitude of lactate enrichment of *F. varium* in one frequently used sample (KSU-Cu), suggesting repeated freeze/thaw cycles were negatively impacting *Fusobacterium* viability.

### *F. varium* grows in selective media used in prior culture methods to quantify *F. necrophorum*

We further evaluated the ability of *F. varium* and *F. necrophorum* ruminal isolates to grow on lactate in two different growth media: Medium II [22] supplemented with 50 mM lactate and modified lactate medium (MLM), which contains lactate as the major carbon source and antibiotics putatively selective for *F. necrophorum*. MLM was previously used for MPN enumeration of *F. necrophorum* in ruminal samples in conjunction with an indole assay [20]. Although *F. varium* growth was not enhanced by lactate supplementation when grown in Medium II, it did display increased maximum cell densities in the presence of lysine and maltose (data not shown). *F. varium* exhibited moderate growth in MLM (Fig.3), suggesting that this species may have been overlooked in previous enumeration estimates. The three *F. varium* strains screened for growth on MLM achieved about 43% of the optical density of the three *F. necrophorum* strains (p-value < 0.05).

**Figure 3.**
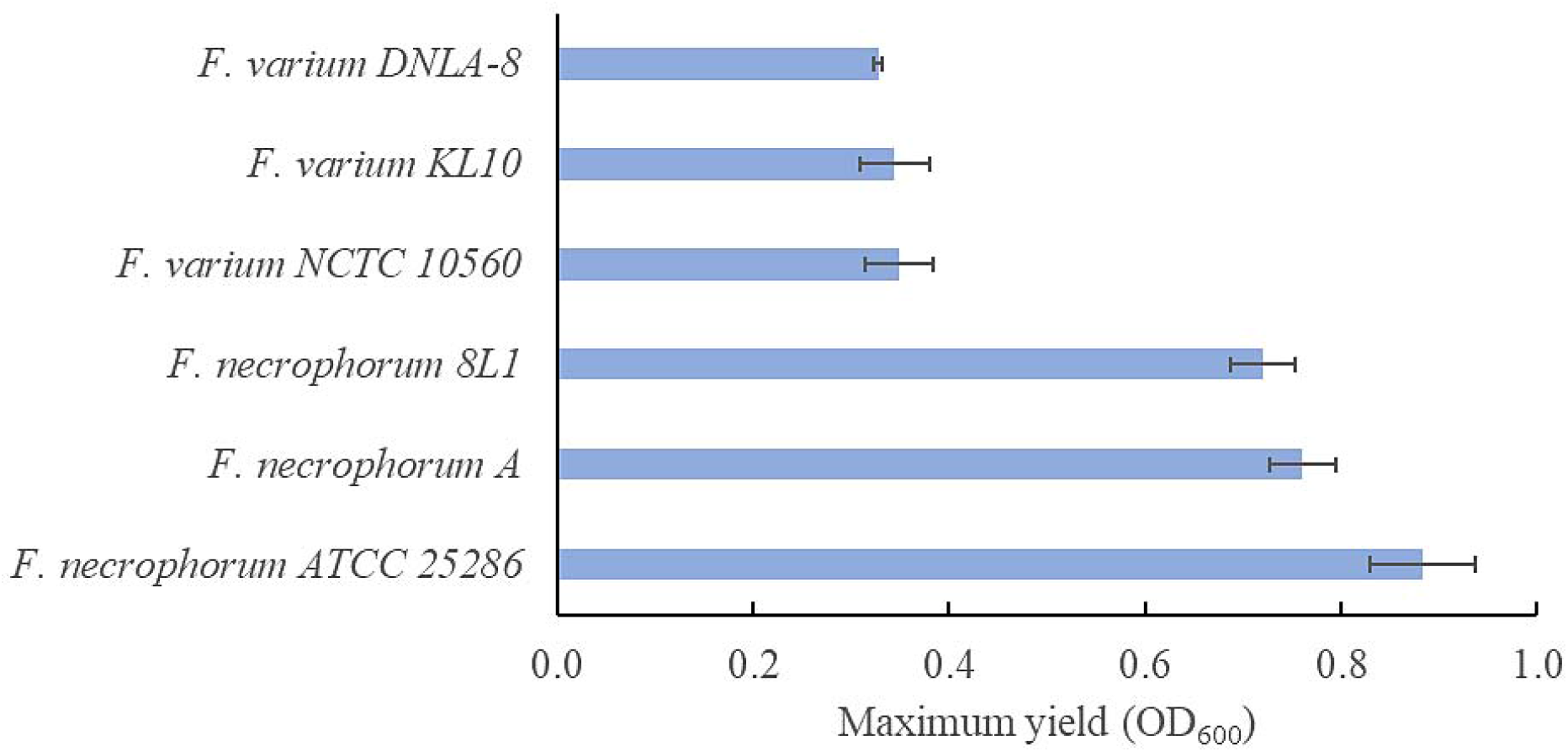
*F. varium* isolates and reference strains grow on a modified lactate medium (MLM) selective for *F. necrophorum*, suggesting that previous culture-based counts of *F. necrophorum* in cow rumen samples may be inflated. Growth observed via OD_600_ measurements of several *F. necrophorum* and *F. varium* strains in a selective modified lactate medium (MLM). The maximum OD_600_ reached by *F. varium* under these conditions was about 43% of that reached by the *F. necrophorum* strains (p-value < 10^−7^).

### Both reference and newly isolated strains of *F. varium* were resistant to common antibiotic feed additives tylosin and monensin

The susceptibility of both reference strains and our new ruminal isolates of *F. varium* and *F. necrophorum* to the macrolide antibiotic tylosin and ionophore monensin were investigated at levels relevant to ruminal concentrations [34-36]. Of the three strains of *F. varium* tested, none were substantially inhibited by tylosin, with the only statistically significant reduction in growth (13%) observed in *F. varium* DNLA-8 (Fig. 4). The three *F. necrophorum* strains tested were all inhibited over 67%, with *F. necrophorum* 8L1 showing a 92% reduction in growth while strain A was most susceptible (94% inhibition).

**Figure 4.**
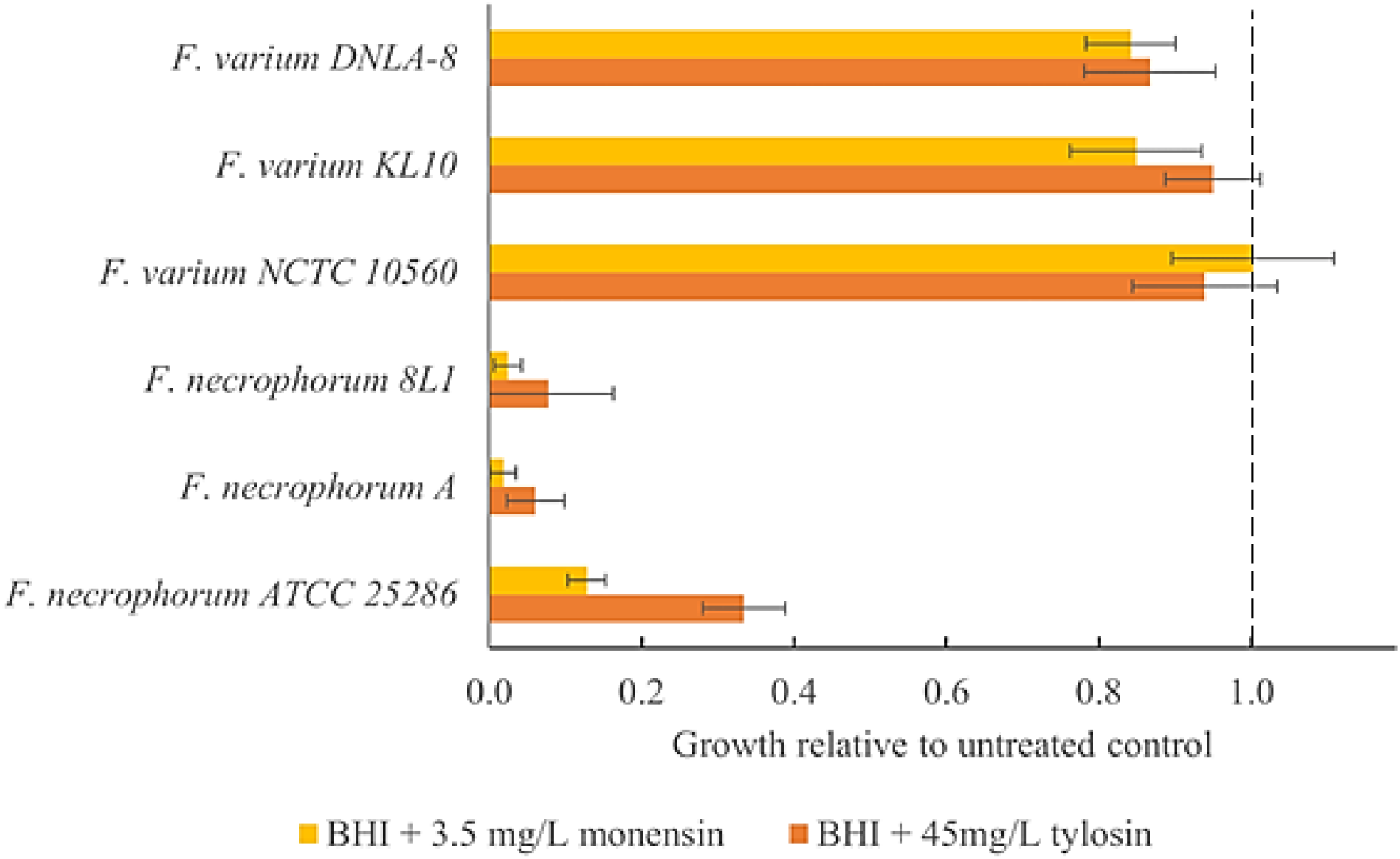
*F. varium* is resistant to both tylosin and monensin. Two reference strains and four *Fusobacterium* ruminal isolates were tested for their susceptibility to two commonly fed antibiotics. Growth (OD_600_) relative to control was strongly inhibited for the three tested *F. necrophorum* strains exposed to 45mg/L tylosin-tartrate or 3.5 mg/L monensin sodium sulfate. In contrast, the *F. varium* strains showed little to no response to these common bovine antibiotics.

Growth of the three *F. necrophorum* strains was reduced more than 87% with the addition of monensin sodium sulfate to the anaerobic BHI medium, while *F. varium* showed little to no response (Fig. 4). Both *F. necrophorum* A and 8L1 showed a 98% reduction in growth, while strain ATCC 25286 was inhibited by 87%. Of the *F. varium* tested, monensin in BHI inhibited growth by 15% in the bovine isolates (p-values < 0.05). Reference strain *F. varium* NCTC 10560 was not significantly inhibited by monensin. When grown in a medium with only yeast extract and casein pancreatic digest as carbon sources (PY), the addition of monensin suppressed *F. necrophorum* to a lesser extent (Fig. S1). *F. necrophorum* 8L1 showed a 66% reduction in growth, while reference strain ATCC 25286 and isolate A both showed a 44% reduction. Growth of the *F. varium* strains was not significantly different from the controls. Overall, our findings suggest that the effect of tylosin and monensin on *F. varium* is negligible, while *F. necrophorum* is generally more susceptible and displays more strain- and context-dependent variability.

### Genome sequencing of ruminal *F. varium* isolates revealed genes associated with virulence

To better understand the potential ecological role that *F. varium* may be playing in the bovine rumen, two *F. varium* ruminal isolates were selected for genome sequencing on the basis of their observed differences in metabolic capacity. *F. varium* 1701-2 displayed strong growth on maltose relative to other *F. varium* strains, while *F. varium* KL10 was found to be highly enriched in PY media controls without lactate. The assembly for *F. varium* KL10 yielded three contigs, totaling 3.36 Mb, while *F. varium* 1701-2 produced two contigs totaling 3.44 Mb. These genome sizes are consistent with sizes reported for the three complete genomes of human isolates deposited in NCBI. The GC content of *F. varium* KL10 and 1701-2, were 29.3 and 29.5 percent, respectively, which is consistent along species lines with the deposited strains *F. varium* NCTC 10560 (29.3%) and *F. varium* Fv113_g1 (29.2%) [40]. The average nucleotide identities (ANI) [41] between both *F. varium* strains and *F. varium* NCTC 10650 were over 98%. The ANI with Fv113_g1. was about 90%, though considering the uncertainty of its taxonomy and similarities to *F. ulcerans* [40], this was not altogether unexpected.

Preliminary genome analysis of the two ruminal *F. varium* isolates using the RAST annotation pipeline [28-30] identified virulence-associated subsystems in both as well as in the previously sequenced human isolates, a summary of which is included in Supplementary Table 1. RAST identified a slightly higher percentage of Virulence, Disease and Defense genes (e.g. metal resistance genes and multidrug efflux pumps) in the ruminal isolates than in both *F. necrophorum* ATCC 25286 [42] and *F. varium* NCTC 10560. However, about 75-80% of the ORFs in each RAST analysis were assigned to no subsystem, suggesting a more extensive analysis of virulence-related genes is needed to assess pathogenic potential. Both *F. varium* isolates contained sequences similar to two-partner secretion systems (TPS), a widespread collection of large virulence exoproteins [43]. The two TPS found in the isolates had the nearest similarity to genes associated with Type V secretion systems, which are known virulence factors within the *Fusobacterium* genus [44]. One TPS was similar to members of the channel-forming TpsB family and the other is similar to members of the ShlA/HecA/FhaA family which are associated with heme utilization or adhesion. Further investigation using the InterPro Database [45, 46], FusoPortal [47, 48], and NCBI databases [49] found sequence similarities to virulence genes found in other *Fusobacterium* species and strains, such as autotransporters, antibiotic resistance genes, and other “active invader” associated proteins [44, 50]. Additionally, strains KL10 and 1701-2 contained genes identified by RAST as similar to the *Mycobacterium* virulence operon possibly involved in quinolinate biosynthesis, which is also found in the human isolate, *F. varium* NCTC 10560.

Among the genes related to lactate metabolism, both isolates contained both D- and L-lactate dehydrogenases, which were located near FMN-dependent ABC transporters. Additionally, the genomes of both isolates contained sequences similar to a lactate utilization gene identified within the deposited strain *F varium* NCTC 10560 and portions with similarity to lactate dehydrogenases found within *F. necrophorum but* lacked the lactate permeases found in other *Fusobacterium* genomes. Among genes related to lysine metabolism, neither isolate contained sequences that matched the lysine permease found in *F. necrophorum, F. nucleatum*, or *F. varium* Fv113_g1, but all contained genes related to lysine degradation.

## DISCUSSION

Overall, our results provide definitive proof that *F. varium* is both widespread and a predominant member of its genus in cattle rumen. Indeed, using enrichment or selective agar, we identified *F. varium* in every sample tested, suggesting it is ubiquitous and indigenous to the bovine rumen. The finding that the most common *Fusobacterium* species in the bovine rumen is often *F. varium*, and that a lactate- and antibiotic-supplemented medium tailored to enrich for *F. necrophorum* fortuitously enriched *F. varium* has implications for past and future interventional studies focused on ruminal acidosis and bovine liver abscess prevention. Importantly, it is likely that previous studies that relied on MPN-indole assays to quantify *F. necrophorum* unintentionally reported inflated cell densities due to the overlooked presence of *F. varium* (which can also produce indole [51, 52]) and reinterpretation of those is warranted. For example, interventions proven effective against *F. necrophorum* in vitro using pure cultures (e.g., limonene [11]) may have been determined to be ineffective during in vivo studies due to the presence of *F. varium*. In contrast, we used a combination of culture-dependent and independent methods that facilitated species-level taxonomic classification, and subsequently sequenced the genomes of two bovine isolates. We found bovine *F. varium* isolates to be highly resistant to antibiotics to which *F. necrophorum* is susceptible, highlighting the potential for differential responses to various treatment modalities.

Beyond the impact on prior studies, it is important to understand the role of *F. varium* in the bovine rumen and whether or not it contributes to ruminal fermentation and LA occurrence and progression, or other disease processes. For several decades, ruminal *F. necrophorum* has been considered the primary etiological agent of liver abscesses, but recent studies have challenged this to varying extents, with data suggesting that Bacteroidetes and Proteobacteria may play a larger role than previously believed and that some LA communities may originate from the hindgut [53, 54]. Since most prior studies were conducted at genus-level taxonomic resolution (e.g., using 16S rRNA sequencing on an Illumina platform), it is currently unclear what role different *Fusobacterium* species may play in liver abscess formation and/or severity, and it cannot be ruled out that *F. varium* may play a direct or indirect role in LA formation. With in-feed tylosin, 12 - 18% of feedlot cattle still develop LAs [14], compared to 30 to 45% when the antibiotic is not fed [7]. An *F. varium* isolate from a respiratory tract abscess of a white-tailed deer was previously reported to be resistant to tylosin [55], and our results corroborate this finding, with all of our bovine isolates displaying almost complete resistance at concentrations several-fold higher than that achieved in practice. Thus, current LA management practices would be ineffective against the bovine *F. varium* strains we isolated from feedlot cattle ruminal fluid.

Feed efficiency, ruminal acidosis and liver abscessation are closely entwined processes that have been studied for decades [56-58], though investigation to elucidate clear causality and mechanistic understanding of ruminal metabolism and microbial community structure is ongoing [59, 60]. *F. necrophorum* has been described as a hyper-ammonia-producing species [61], and is believed to negatively impact feed efficiency [62], possibly through lysine fermentation. A dose-dependent correlation has been observed between liver abscessation and lysine supplementation during the finishing of beef steers [63]. Monensin was reported to reduce the uptake of lysine by bacteria, though *F. necrophorum* appears to have varying susceptibility that is dependent on lactate and lysine availability [37]. Interestingly, we found *F. varium* growth to be unaffected by monensin. Genomic analysis of both bovine *F. varium* isolates revealed the presence of genes encoding enzymes involved in lysine fermentation, suggesting a possible negative impact on post-ruminal lysine availability, which in lactating dairy cows is a limiting amino acid for milk production. Lysine degradation has also been documented in *F. varium* in prior studies [64], though we found no explicit lysine-specific permeases within the *F. varium* isolates examined. In conjunction with the greater abundance of this organism in the rumen samples analyzed, this finding may have relevance for strategies that use monensin to increase feed efficiency, reduce ruminal acidosis, or control bloat in cattle [65, 66].

*F. necrophorum* has also been reported to be the only *Fusobacterium* species able to utilize lactate as a carbon and energy source [20, 51], though there has been speculation that small amounts of lactate metabolism may occur in other *Fusobacterium* species in the presence of amino acids, as well as conflicting reports of *F. varium* enhanced growth in media supplemented with lactic acid [17, 67, 68]. Though we did not find unequivocal evidence that *F. varium* is utilizing lactate for energy production, all *F. varium* genomes analyzed carry lactate dehydrogenases, and we routinely observed robust increases in cell densities (up to 4-log) in media containing lactate as the primary energy source. While no explicit lactate permeases were detected in our preliminary analysis, this does not necessarily exclude their presence [69-71]. Note that we routinely observed very substantial enrichment of *F. varium* using only lactate amendment. In general, our data suggest a potentially important unelucidated role within established acidosis-rumenitis-liver abscess theory. Overlapping metabolic capacity is also important to consider as it is a major determining factor for interactions between *F. necrophorum* and *F. varium*, which could help inform efforts to selectively modulate either species. Overall, our results demonstrate that the current practices used to control *F. necrophorum* are not effective against *F. varium* and could be inadvertently enhancing its growth due to a reduction in competition.

It is unknown whether *F. varium* dominance of the ruminal *Fusobacterium* subpopulation is due to changes in ruminant production practices [72, 73], recent acquisition and proliferation within US herds, or whether it has simply been overlooked. However, the latter possibility (overlooked predominance) is not altogether surprising – typically, the relative abundance of the *Fusobacterium* genus in the rumen community is low, around 0.002% [38] and surveys of these communities are rarely conducted beyond the genus level. However, despite its low relative abundance, interest in the *Fusobacterium* genus has rapidly grown due to its increasing association with numerous human and animal diseases and infections [44, 74]. Though little has been reported of its presence in cattle, *F. varium* has been associated with ulcerative colitis in Japanese patients but has not been as frequently reported in North America or Europe [75, 76]. Notably, however, a recent study found a rapid increase in the proportion of *Fusobacterium* infections in Korean patients caused by *F. varium* since 2016, referring to the species as an emerging pathogen: *F. varium* was shown to cause more severe infections than *F. nucleatum* and *F. necrophorum*, demonstrated by high rates of in-hospital mortality (12.5%) and ICU admission (26.8%) as well as acute kidney injury (14.3%) [21]. Thus, it is possible that *F. varium* is also an emerging cattle pathogen, though more work is needed to justify concern.

While the current understanding of *Fusobacterium* virulence mechanisms remains limited, species are broadly categorized as either actively invasive, passively invasive (requiring a compromised epithelial barrier for entry) or having unknown invasive potential [44]. Notably, *F. necrophorum* is considered passively invasive, while *F. varium* is considered actively invasive [77]. Compared to passive invaders, active invaders generally have larger genomes encoding more genes related to host cell invasion. Considering the lack of complete genomes of bovine-isolated *F. varium*, we sequenced the genomes of two ruminal *F. varium* isolates (strains FV-KL10 and FV-1701-2). The two isolates had a high similarity (> 98%) to *F. varium* NCTC 10560 and ATCC 27725 but a lower similarity (90%) to the other human isolate with a complete genome, Fv113_g1. Interestingly, investigation into the substantially expanded genome of Fv113_g1 includes mobile genetic elements related to lactate utilization and virulence. Notably, we have observed considerable variability in the metabolic capacity of our bovine *F. varium* isolates, highlighting the potential for functionally-distinct bovine strains. Thus, a more thorough investigation of bovine *F. varium* genomes may be warranted, particularly to assess whether such mobile genetic elements are present as they could be related to the recent emergence of *F. varium* as a human pathogen in several Asian countries [21, 78].

The pathogenic mechanism of *F. necrophorum* is primarily ascribed to a unique leukotoxin [79], the sequence of which we did not detect within any of our *F. varium* isolates. However, both bovine *F. varium* isolates contained sequences with high similarities to the FadA homologues found in the known human pathogens *F. varium* NCTC 10560 and *F. varium* Fv113_g1. FadA is a small adhesin which multimerizes on the cell surface and has been best characterized in *F. nucleatum* where it was shown to be involved in β-catenin signaling and regulating the inflammatory and oncogenic responses in a colorectal cancer mice model [79-81]. Phylogenetic analysis has identified a family of FadA-like adhesin genes are present across *Fusobacterium* species, with the invasive strains containing a higher number and diversity [44]. Autotransporter proteins and sequences similar to type V secretion system (T5SS) were also detected in our cursory genomic analyses, as well as multidrug efflux pumps, beta-lactamases, fluoroquinolone resistance genes, and putative internalin genes, which are involved in invasion and intracellular resistance [82]. RAST also predicted genes related to quinolinate synthesis, which is a metabolite in the tryptophan catabolism pathway but also a potent neurotoxin [83].

A lack of genetic tools for manipulating *Fusobacterium* genomes has hindered research to determine the role of many predicted virulence factors in this genus. Thus, while *F. varium* encodes a larger number of putative virulence factors relative to *F. necrophorum*, its exact role has not been delineated at this time, highlighting the need for further research.

## CONCLUSIONS

Though historically overlooked, our results demonstrate that *F. varium* is a widespread bovine ruminal resident with an unknown ecological role. However, recent trends in the species’ proliferation and association with various human and animal diseases and infections is a potential concern. That bovine isolates of *F. varium* contain genes for cellular invasion and pathogenesis similar to their human counterparts may have important implications for the cattle industry in regard to liver abscess prevention. As such, its presence within the bovine rumen is an important finding and its role within the ruminal community requires further investigation. Considering recent interest in the virulence of members of the *Fusobacterium* species and the trend of overlooking *F. varium* as an important player in the ruminal compartment, this research brings timely attention to this species, and encourages revisiting the etiology of bovine ruminal ulceration and liver abscess formation.

## Acknowledgements

This work was funded by Small Business Innovation Research (SBIR) grants from the National Science Foundation (2025980) and the United States Department of Agriculture (2020-00650).

