## Supplemental Information for "Unexpected finding of *Fusobacterium varium* abundance in cattle rumen: implications for liver abscess interventions"

**Running Head: *F. varium* highly abundant in bovine rumen**

**Table S1. . Subsystems assigned by the RAST platform of several deposited strains and two new bovine *F. varium* isolates.**

| Subsystem Feature Counts | *F. necrophorum* ATCC 25286 | *F. varium* NCTC 10560 | *F. varium* Fv133_g1 | *F. varium* KL10 | *F. varium* 1701-2 |
| --- | --- | --- | --- | --- | --- |
| Cofactors, Vitamins, Prosthetic Groups, Pigments | **4.16%** | **3.24%** | **2.67%** | **3.31%** | **3.21%** |
| Cell Wall and Capsule | **1.61%** | **1.65%** | **1.14%** | **1.26%** | **1.48%** |
| Virulence, Disease and Defense | **1.17%** | **1.62%** | **1.55%** | **2.08%** | **1.86%** |
| Potassium metabolism | **0.08%** | **0.17%** | **0.21%** | **0.15%** | **0.15%** |
| Miscellaneous | **0.32%** | **0.20%** | **0.16%** | **0.18%** | **0.17%** |
| Phages, Prophages, Transposable elements, Plasmids | **0.12%** | **0.17%** | **0.21%** | **0.37%** | **0.13%** |
| Membrane Transport | **2.02%** | **1.25%** | **0.73%** | **1.49%** | **1.43%** |
| Iron acquisition and metabolism | **0.61%** | **0.14%** | **0.14%** | **0.34%** | **0.21%** |
| RNA Metabolism | **1.13%** | **1.08%** | **1.12%** | **1.26%** | **1.13%** |
| Nucleosides and Nucleotides | **2.38%** | **2.36%** | **2.09%** | **2.84%** | **2.44%** |
| Protein Metabolism | **4.97%** | **3.27%** | **3.07%** | **4.02%** | **3.78%** |
| Regulation and Cell signaling | **0.52%** | **0.26%** | **0.19%** | **0.37%** | **0.24%** |
| DNA Metabolism | **2.70%** | **2.36%** | **1.96%** | **2.12%** | **2.73%** |
| Fatty Acids, Lipids, and Isoprenoids | **1.57%** | **1.28%** | **1.01%** | **1.34%** | **1.09%** |
| Dormancy and Sporulation | **0.08%** | **0.20%** | **0.21%** | **0.23%** | **0.26%** |
| Respiration | **1.09%** | **0.97%** | **0.71%** | **1.12%** | **0.83%** |
| Stress Response | **0.89%** | **0.74%** | **0.76%** | **0.89%** | **0.75%** |
| Metabolism of Aromatic Compounds | **0.08%** | **0.03%** | **0.13%** | **0.05%** | **0.06%** |
| Amino Acids and Derivatives | **3.67%** | **5.85%** | **5.36%** | **6.67%** | **6.69%** |
| Sulfur Metabolism | **0.08%** | **0.06%** | **0.11%** | **0.06%** | **0.08%** |
| Phosphorus Metabolism | **0.00%** | **0.63%** | **0.43%** | **0.65%** | **0.86%** |
| Carbohydrates | **3.79%** | **5.40%** | **4.79%** | **6.53%** | **5.94%** |
| Subsystem coverage | **25.47%** | **23.42%** | **20.18%** | **25.92%** | **24.92%** |
| Not in Subsystem | **74.53%** | **76.58%** | **79.81%** | **74.08%** | **75.08%** |

Maximum yield (OD600)

|  | PY | PY + 3.5 mg/L monensin sodium sulfate |
| --- | --- | --- |
| F. necrophorum ATCC 25268 | 0.968175 | 0.540875 |
| F. necrophorum A | 0.9011 | 0.507125 |
| F. necrophorum 8L1 | 0.87105 | 0.2967 |
| F. varium NCTC 10650 | 0.536042 | 0.644175 |
| F. varium KL10 | 0.600508 | 0.635025 |
| F. varium DNLA-8 | 0.567742 | 0.607917 |

**Figure S1. Efficacy of monensin on F. necrophorum is context dependent, though F. varium remains unaffected.** In a context with a lesser amino acid concentration (PY medium), the reduction in growth of *F. necrophorum* is less than in BHI when challenged with monensin. The effect of monensin in PY versus untreated controls on *F. varium* was not statistically significant.
